# Renal Angiotensinogen is Predominantly Liver-Derived in Nonhuman Primates

**DOI:** 10.1101/2021.05.18.444555

**Authors:** Masayoshi Kukida, Lei Cai, Dien Ye, Hisashi Sawada, Yuriko Katsumata, Michael K. Franklin, Peter I. Hecker, Kenneth S. Campbell, A.H. Jan Danser, Adam E. Mullick, Alan Daugherty, Ryan E. Temel, Hong S. Lu

## Abstract

AGT (Angiotensinogen) is the unique substrate of the renin-angiotensin system. Liver is the primary source of circulating AGT. The present study determined whether hepatocyte-derived AGT regulates renal AGT accumulation by injecting ASO (antisense oligonucleotides) targeting hepatocyte-derived AGT (GalNAc AGT ASO) into female cynomolgus monkeys. Hepatocyte-specific inhibition of AGT led to profound reductions of plasma AGT concentrations. AGT protein in S1 and S2 of renal proximal tubules was greatly diminished by GalNAc AGT ASO. Given the similarity between nonhuman primates and human, our findings support the notion that renal AGT is predominantly derived from liver, and liver regulates renal angiotensin II production in humans.

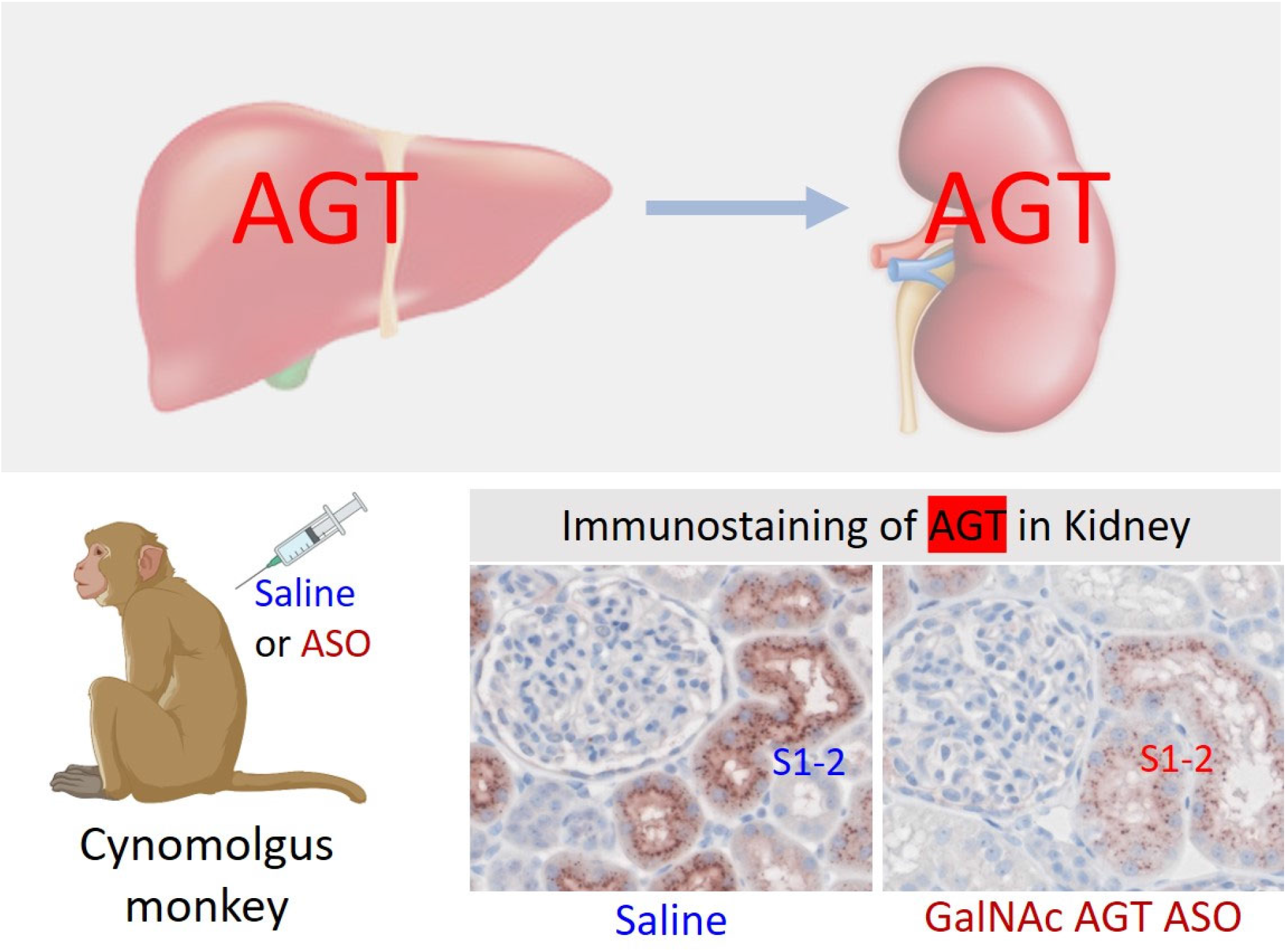

---

AGT (Angiotensinogen) is the unique substrate of the RAS (renin-angiotensin system). While many cells synthesize AGT, studies in mice have demonstrated that plasma AGT is predominantly liver-derived.^1, 2^ Indeed, either pharmacological inhibition or genetic deficiency of hepatocyte-derived AGT reduces blood pressure and atherosclerosis in mice.^3–5^ Locally synthesized AGT in the kidney may contribute to renal Ang (angiotensin) II generation, possibly in a blood pressure-independent manner, although the literature is inconsistent.^6, 7^ Based on this concept, AGT measurements in urine or renal biopsies are used often as an independent marker of the renal RAS activation in humans. However, it remains unclear whether renal Ang II generation in humans depends on kidney-derived AGT.

AGT is cleaved by renin into two products: Ang I, which consists of 10 amino acids, and des(Ang I)AGT, which has 443 amino acids in mice and 442 amino acids in humans and NHP (nonhuman primates). Although sequences of AGT vary substantially between mouse and human, this protein is highly conserved in humans and NHP. Plasma total AGT concentrations were 3-4 μg/mL in mice, 15-41 μg/mL in humans, and 11-20 μg/mL in cynomolgus monkeys (**Figure 1**). Plasma AGT was predominantly present as des(AngI)AGT in mice (~92%) in contrast to both humans and cynomolgus monkeys (< 40% of des(AngI)AGT).

**Figure 1.**
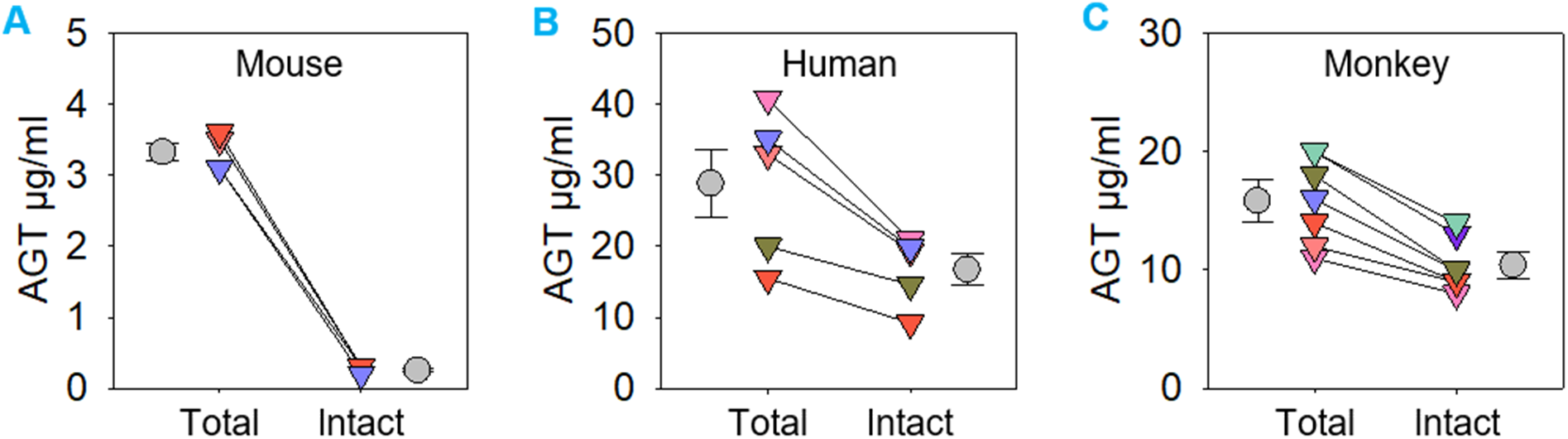
Plasma total AGT and intact AGT concentrations in (A) male C57BL/6 mice (N=4), (B) humans (N=5), and (C) cynomolgus monkeys (N=7) were measured using ELISA kits (mouse ELISA kits: IBL America 27413 for total AGT and 277423 for intact AGT, human and monkey ELISA kits: IBL America 27412 for total AGT and 27742 for intact AGT). Total AGT = Intact AGT + des(Angl)AGT.

Despite differences in plasma AGT, the distribution of AGT protein accumulation within the kidney was comparable among the 3 species. AGT protein accumulation was most abundant in the renal proximal convoluted tubules (S1 and S2 segments), modest in the proximal straight tubules (S3 segment), and not detected in glomeruli and other tubules of the kidneys (**Figure 2**).

**Figure 2.**
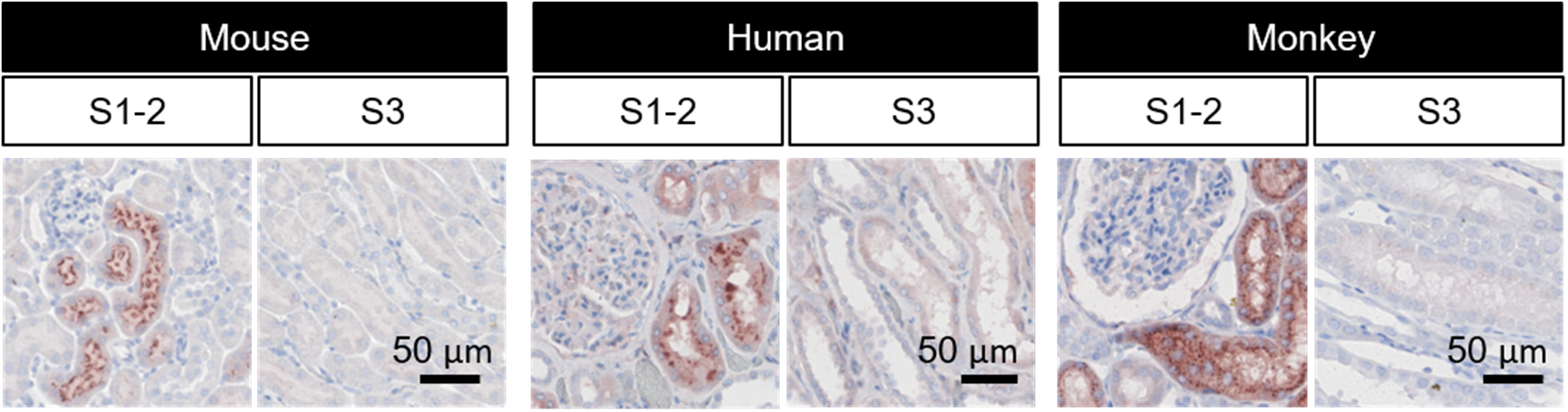
Immunostaining of AGT in kidney sections from mice, humans, and cynomolgus monkeys using a rabbit anti-mouse AGT antibody for mouse (IBL America 28101; orking concentration: 0.3 (jg/ml) and a mouse anti-human AGT antibody for both humans and monkeys (IBL America 10417; working concentration: 1 pg/ml).

Given the similarity between humans and cynomolgus monkeys, findings in the latter likely have greater translational significance, compared to rodent models, in defining the origin of kidney AGT in humans. To determine whether liver-derived AGT contributes to AGT protein accumulated in the kidney of NHP, female cynomolgus monkeys (3-4 years of age) were injected subcutaneously with either saline or ASO (antisense oligonucleotides) targeting liver-derived human AGT (Ionis, GalNAc AGT ASO: conjugated with N-acetyl galactosamine; 2.5 or 10 mg/kg). This human GalNAc AGT ASO has an identical sequence match to AGT mRNA in cynomolgus monkeys. Saline or ASO was injected on day 1 and day 4, and then once weekly for a subsequent 4 weeks. Neither dose of ASO affected body weight or liver and kidney functions.

Both doses of GalNAc AGT ASO reduced plasma AGT concentrations within 1 week by up to 80% (**Figure 3A**). Liver had approximately 160-fold more AGT mRNA abundance than kidney and visceral adipose tissue (**Figure 3B**). Both doses of GalNAc AGT ASO profoundly reduced hepatic mRNA abundance of AGT (**Figure 3C**). The low dose (2.5 mg/kg) of GalNAc AGT ASO did not affect renal AGT mRNA abundance, whereas the high dose (10 mg/kg) reduced renal AGT mRNA abundance (**Figure 4A**). Of note, both doses of GalNAc AGT ASO produced equivalent reductions in both plasma and liver AGT. This illustrates that GalNAc ASO can be detected in kidney and exhibits some activity at sufficiently high doses such as 10 mg/kg in cynomolgus monkeys. Irrespective of renal AGT mRNA, both doses of GalNAc AGT ASO diminished renal AGT protein accumulation to a similar extent (**Figure 4B**). Plasma renin activity was not altered by either dose of GalNAc AGT ASO (**Figure 4C**), consistent with the recent data evaluating IONIS-AGT-L_RX_ in hypertensive patients.7 These data imply that renin upregulation must have matched AGT downregulation to keep angiotensin generation in the normal range.^6^

**Figure 3.**
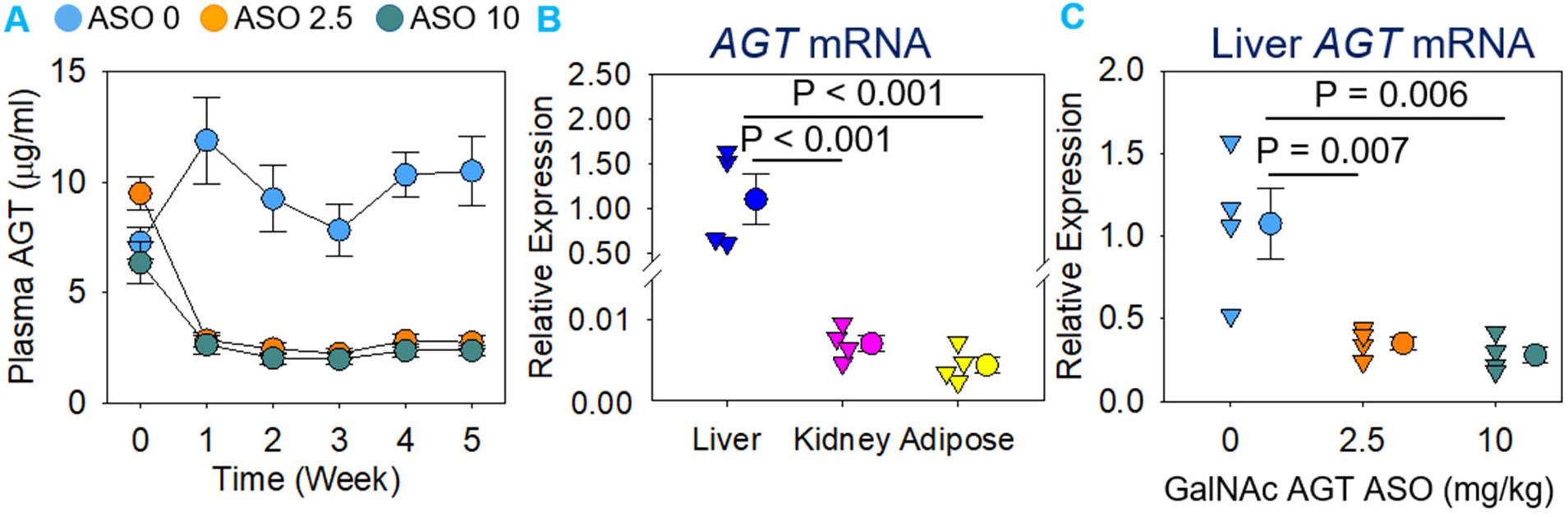
Female cynomolgus monkeys were injected subcutaneously with either saline (indicated as “ASO 0” in the figure) or GalNAc AGT ASO (2.5 or 10 mg/kg; indicated as “ASO 2.5” and “ASO 10”, respectively) for 5 weeks (N = 4/group). (A) Plasma total AGT concentrations were determined by an ELISA kit (IBL America 27412). Data are represented as mean ± SEM. Piecewise linear mixed model with a split point at week 1 was used to compare plasma AGT concentration changes over time among the three groups. P < 0.001 between week 0 and week 1 for both ASO 2.5 and ASO 10, compared to ASO 0, and the differences remained during the study. (B, C) mRNA abundance of AGT was quantified by qPCR. Data were calculated using the AACt method and normalized to the mean of two reference genes: GAPDH (Glyceraldehyde-3-phosphate dehydrogenase) and ZFP91 (Zinc finger protein 91). Data were Iog10 transformed for normality and equal variance tests.

**Figure 4.**
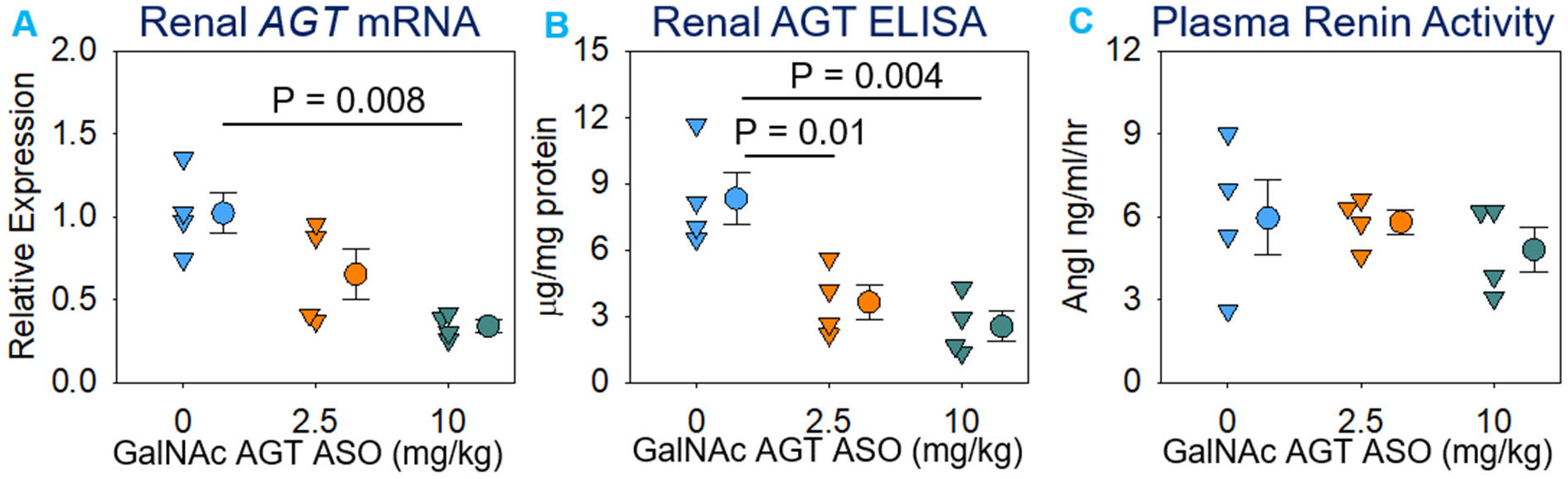
Female cynomolgus monkeys were injected subcutaneously with either saline or (2.5 or 10 mg/kg, respectively) for 5 weeks (N = 4/group). (A) mRNA AGT was quantified by qPCR. (B) Renal AGT protein concentrations were an ELISA kit (IBL America 27412) and normalized by total protein Plasma renin activity was determined using an angiotensin I ELISA kit (IBL Since all data passed normality and equal variance test, all data were ?way analysis of variance with the Holm-Sidak method.

As shown by immunostaining, diminished AGT protein accumulation following dosing with GalNAc AGT ASO was noted in the S1 and S2 segments of renal proximal tubules (**Figure 5**). These findings support the notion that liver supplies the bulk of AGT protein to the kidney in NHP, independent of the presence of renal AGT mRNA. **Figure 5** also shows the distribution of other RAS components. Renin was observed predominantly in juxtaglomerular cells, and ACE (angiotensin-converting enzyme) and ACE2 were present in all 3 segments of the proximal tubules, being most abundant in the S3 portion. GalNAc AGT ASO did not change the renal distribution of these enzymes.

**Figure 5.**
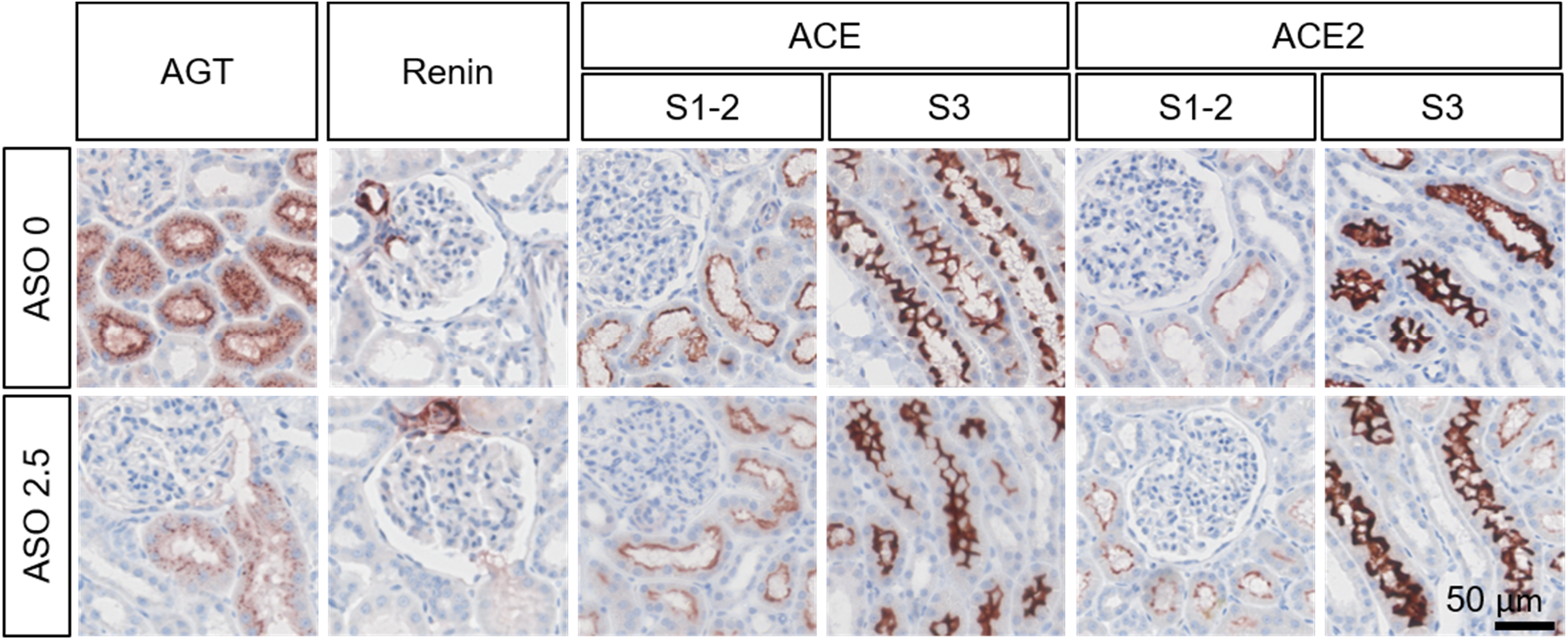
Immunostaining of AGT, renin, ACE, and ACE2 in kidney sections. Antibody information: mouse anti-human AGT (IBL America 10417; 1 |jg/ml), rabbit anti-human renin (Sigma HPA005131; 1 pg/ml), rabbit anti-human ACE (Sigma HPA029298; 2 jjg/ml), and rabbit anti ACE2 (Abeam ab108252; 0.3 |jg/ml). Mouse anti-human AGT antibody (IBL America 10417) recognizes the C-terminus of AGT, it detects both intact AGT and its cleaved form, des(Angl) AGT. Saline is indicated as "ASO 0" and GalNAc AGT ASO 2.5 mg/kg is indicated as "ASO 2.5".

In conclusion, the liver is the major source of AGT in kidneys of cynomolgus monkeys. Hepatic AGT accumulation in the S1 and S2 segments of the proximal tubules coincides with the observation in humans that tubular reabsorption via megalin, an endocytic receptor on the proximal tubules, is the main determinant of urinary AGT.^1, 8^ Taken together, these data are consistent with renal AGT originating predominantly in the liver. This implies that renal Ang II production in NHP and humans relies on hepatic AGT, and that the concept that AGT in urine or renal biopsies reflects an independent renal RAS needs to be reconsidered.

## Acknowledgment

We thank CCTS Biospecimens Core (Supported by UL1TR001998) at the University of Kentucky for providing human samples.

## Abbreviations

AGT: Angiotensinogen
Ang: Angiotensin
RAS: Renin angiotensin system
ASO: Antisense oligonucleotides
NHP: Nonhuman primates
ACE: Angiotensin-converting enzyme

## MATERIALS AND METHODS

### Animals

C57BL/6J (#000664) mice were purchased from The Jackson Laboratory (Bar Harbor, ME, USA). All mice were maintained in a barrier facility on a light: dark cycle of 14: 10 hours, and fed a normal mouse laboratory diet. Blood samples were collected by right ventricular punctures at termination.

Adult male and female cynomolgus monkeys (Macaca fascicularis) were purchased from DSP Research Services and used for the non-human primate studies. All monkeys were housed in climate-controlled conditions with 12-hour light/dark cycles, and fed Teklad Global 20% Protein Primate Diet. After overnight fast, monkeys were anesthetized with ketamine (5 mg/kg) and dexmedetomidine (0.0075 - 0.015 mg/kg), and blood was collected from the femoral vein. Body weights were measured in unconscious state. For GalNAc AGT ASO studies, female monkeys were injected subcutaneously with either vehicle (saline, USP grade) or GalNAc AGT ASO (2.5 or 10 mg/kg). Saline or ASO was injected on day 1 and day 4, and then once weekly for the subsequent 4 weeks.

All experiments performed were approved by the University of Kentucky Institutional Animal Care and Use Committee.

### Plasma profiles

Blood samples from mice and monkeys were collected into tubes with EDTA (final concentration: 1.8 mg/ml). Plasma was collected by centrifugation at 3,000 rcf for 10 minutes, 4 °C, and frozen at −80°C until analysis.

Male human plasma samples were kindly provided by the Center for Clinical and Translational Science Biospecomens Core, University of Kentucky.

Plasma total AGT and intact AGT concentrations were measured by using ELISA kits (mouse ELISA kits: IBL America 27413 for total AGT and 277423 for intact AGT, human and monkey ELISA kits: IBL America 27412 for total AGT and 27742 for intact AGT).

Plasma renin activity in cynomolgus monkeys was measured using an ELISA kit (IBL America IB57101) following the protocol provided from the manufacturer. Briefly, monkey plasma samples were incubated in the assay buffer including a generation buffer and PMSF (phenylmethylsulfonyl fluoride) without exogenous AGT at 37 °C for 90 minutes. The reaction was terminated by placing samples on ice for 5 minutes. AngI generated in each sample was quantified by the ELISA.

### AGT concentrations in kidney

To quantify AGT protein in kidney, we extracted proteins from the outer cortex of kidney (did not include medulla) using a T-per buffer (ThermoFisher Scientific 78510) including with a protease inhibitor cocktail (Cell Signaling Technology 5871S) and measured renal AGT concentrations using the human AGT ELISA kit (IBL America 27412) that was also used for plasma AGT measurements. Total AGT concentration in each sample was normalized by total protein concentrations measured using a DC™ protein assay kit (Bio-Rad 5000112).

### Immunostaining

Isolated whole kidneys from mice and monkeys were fixed in either 10% neutrally buffered formalin at room temperature or cold paraformaldehyde (PFA, 4% wt/vol) solution at 4°C, overnight. Following the tissue-fixation, kidneys were embedded in paraffin, and sectioned 4 μm thick. Human kidney paraffin sections were purchased from the Zyagen Labs (Zyagen, HP901).

After antigen retrieval with citrate buffer (pH 6.0) and blocking, kidney sections were incubated with primary antibodies at 37 °C for 15 minutes or 4°C overnight (Table I), and subsequently incubated with HRP conjugated secondary antibody polymer detection kit (Table II) at room temperature for 30 min. AEC substrate kit (Vector, SK-4200) was used to develop red color for positive staining. After counter staining with hematoxylin (Mayer’s Hematoxylin solution, Electron microscopy sciences 26043-06), stained slides were mounted with glycerol gelatin. Images were captured using Axio scan (ZEISS Axio Scan.Z1, Zeiss). Negative controls included nonimmune Isotype-matched IgG to replace the primary antibodies, omission of primary antibody control, and no primary and secondary antibody control.

**Table I.**
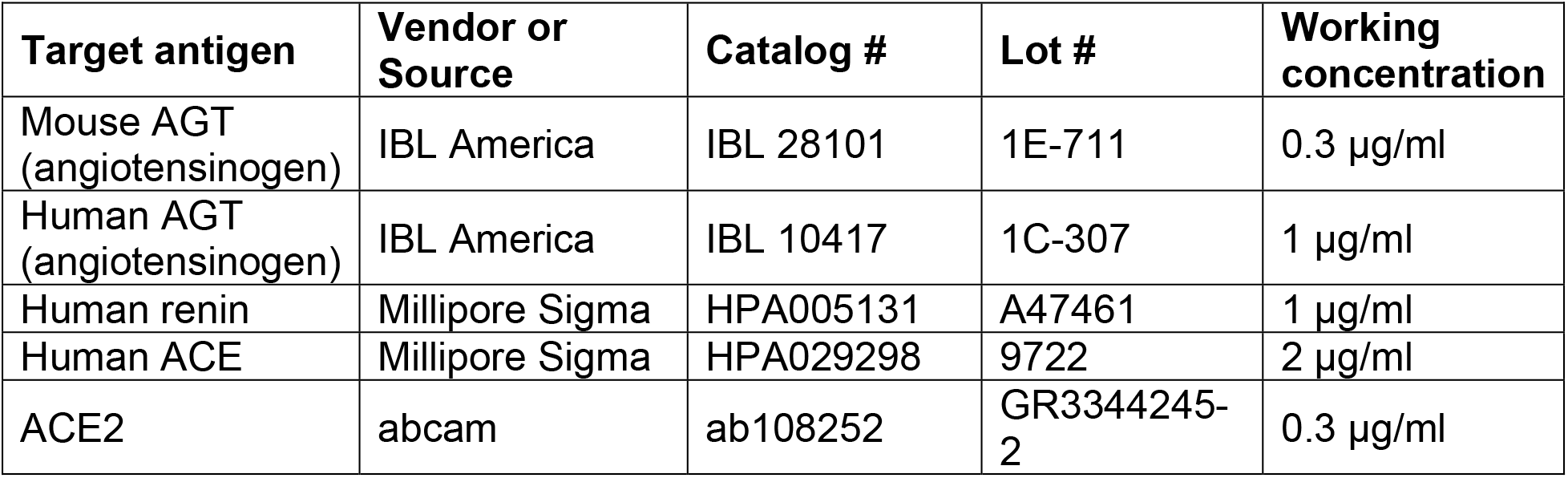
Primary antibodies.

| Target antigen | Vendor or Source | Catalog # | Lot # | Working concentration |
| --- | --- | --- | --- | --- |
| Mouse AGT (angiotensinogen) | IBL America | IBL 28101 | 1E-711 | 0.3 µg/ml |
| Human AGT (angiotensinogen) | IBL America | IBL 10417 | 1C-307 | 1 µg/ml |
| Human renin | Millipore Sigma | HPA005131 | A47461 | 1 µg/ml |
| Human ACE | Millipore Sigma | HPA029298 | 9722 | 2 µg/ml |
| ACE2 | abcam | ab108252 | GR3344245-2 | 0.3 µg/ml |

**Table II.** Detection Kit.

| Kit | Vendor or Source | Catalog # | Lot # |
| --- | --- | --- | --- |
| ImmPRESS® HRP goat anti-rabbit IgG polymer detection kit | Vector | MP-7451 | ZG0629 |
| ImmPRESS® HRP horse anti-mouse IgG, rat adsorbed polymer detection kit | Vector | MP-7422 | ZG0520 |

### Quantitative PCR

Total RNA of the renal cortex in cynomolgus monkeys was extracted using Maxwell RSC simplyRNA tissue kit (Promega, AS1340) by an automated Maxwell RSC Instrument (Promega). To quantify the abundance of mRNA, total RNA was reversely transcribed with an iScript™ cDNA Synthesis kit (Bio-Rad, 170-8891), and quantitative PCR (qPCR) was performed to quantify mRNA abundance using a SsoFast™ EvaGreen^®^ Supermix kit (Bio-Rad, 172-5204,) on a Bio-Rad CFX96 cycler. Data were analyzed using the ΔΔCt method and normalized to the mean of two reference genes: GAPDH (Glyceraldehyde-3-phosphate dehydrogenase) and ZFP91 (Zinc finger protein 91).

## Statistical analyses

The data were represented as individual data points and mean ± SEM. SigmaPlot version 14.0 (Systat Software Inc., San Jose, CA) was used for statistical analyses for Figures 3B, 3C, 4B, and 4C. Normality and homogeneity were tested in all data by Shapiro-Wilk and Brown-Forsythe tests, respectively. Data for AGT mRNA abundance in tissues and liver were log10 transformed to pass normality and equal variance tests. Since the data passed both tests, all data were analyzed using one-way ANOVA with Holm-Sidak method. For plasma AGT concentrations (Figure 3A), piecewise linear mixed model with a split point at Week 1 was performed to compare plasma AGT changes over time among the 3 groups using the “nlme” R package. The estimates of the fixed effect parameters were weighted by the inverse variances at each time point. P < 0.05 was considered statistically significant.

